# The evolution of genetic diversity in changing environments

**DOI:** 10.1101/004440

**Authors:** Oana Carja, Uri Liberman, Marcus W. Feldman

## Abstract

The production and maintenance of genetic and phenotypic diversity under temporally fluctuating selection and the signatures of environmental and selective volatility in the patterns of genetic and phenotypic variation have been important areas of focus in population genetics. On one hand, stretches of constant selection pull the genetic makeup of populations towards local fitness optima. On the other, in order to cope with changes in the selection regime, populations may evolve mechanisms that create a diversity of genotypes. By tuning the rates at which variability is produced, such as the rates of recombination, mutation or migration, populations may increase their long-term adaptability. Here we use theoretical models to gain insight into how the rates of these three evolutionary forces are shaped by fluctuating selection. We compare and contrast the evolution of recombination, mutation and migration under similar patterns of environmental change and show that these three sources of phenotypic variation are surprisingly similar in their response to changing selection. We show that knowing the shape, size, variance and asymmetry of environmental runs is essential for accurate prediction of genetic evolutionary dynamics.

## Introduction

Under constant selection, a large haploid population is expected to evolve towards a local fitness maximum. However, in natural populations selection may not be constant over time due, for example, to ecological changes, spatial variability, changes in lifestyle or even shifts in the genetic background (1). Under temporal and spatial heterogeneity in the direction and strength of selection, a population may evolve mechanisms that create and maintain a stable diversity of phenotypes, thus increasing the long-term adaptability of the population (2-7). These mechanisms may include tuning the rates at which genetic variability is produced, such as the rates of recombination, mutation or migration (8).

Understanding how population genomic dynamics are shaped by fluctuating selection has constituted an important component of mathematical evolutionary theory over the past five decades. These studies have addressed such issues as the relationship between volatility in selection and the dynamics of evolution and how this volatility is reflected in the pattern of genotypic frequency variation (9-12).

One important contributor to the pattern of genetic diversity is recombination, which can affect variation by bringing together or breaking apart combinations of alleles. Recombination may accelerate adaptation by expediting the removal of combinations of deleterious alleles from the population, but also slow it down by breaking apart favorable interactions among genes (13-18). The prevalence of recombination in nature has stimulated theoretical efforts to determine the evolutionarily stable recombination rate under a wide variety of modeling assumptions (14-20). Most explanations of the advantage of sex and recombination involve either rebuilding good combinations of alleles from bad ones created by mutation (19-21), or adapting to a changing environment (14, 22, 23).

One of the earliest models for the evolution of recombination in a fluctuating environment is due to Charlesworth (14), who showed that for a diploid genetic system, when the sign of the linkage disequilibrium varies in time, increased recombination may be favored if the period of environmental fluctuation is strictly larger than two. This is because, when the selective environment varies between positive and negative epistasis, the build-up of disequilibrium is often of the opposite sign to the current epistasis. This lag between epistasis and linkage disequilibrium leads to a mismatch between combinations of alleles that are most fit and those that are more common, and recombination should be favored because it breaks apart the currently maladapted allele combinations and combines alleles that might constitute a fitter haplotype in the future (14, 17, 22, 23).

Environmental fluctuations can also affect the evolution of the genomic mutation rate. In bacteria, stress can increase the mutation rate by inducing mutagenic mechanisms such as the SOS transcriptional response (24, 25) or contingency loci (26). Theoretical studies have suggested that mutation rates should evolve in synchrony with the rate of environmental change. Early studies (27-29) showed that when selection fluctuates periodically and symmetrically between two states with different optimal genotypes, the mutation rate between allelic states will evolve to approximately 1/*n*, where *n* is the number of generations between temporal environmental changes. Similar results were presented by Salathe et al. (29), although the stable mutation rate became zero as the variability in temporal fluctuations increased, or if there were asymmetries in selection pressure between the two environmental states.

Variability in selection has also been shown to drive changes in the evolutionarily stable migration rate. When selection is heterogeneous in space but not in time, migrants cannot displace locally adapted individuals and there is selection against migration (30-32). This has been shown in population genetic models of selection in multi-patch environments with patch-dependent selection of alleles (30), and in ecological analyses of the evolution of dispersal (33-36). Migration is suppressed since, in the absence of temporal variation, it is a force that limits the ability of populations to adapt to local environmental conditions (37). With temporal variability in selection however, migration can increase the level of local adaptation (38) and higher rates of migration may be favored (39-41). Since the pattern of migration affects the structure of populations, understanding the evolution of migration is important for theories of speciation, extinction, host-parasite interactions, or multi-level selection.

Here, we use the mathematical framework of modifier theory to study how environmental fluctuation affects the rates of recombination, mutation and migration. A modifier approach to study the evolution of genetic systems was first introduced by Nei (42) and Feldman (50) in models on the evolution of recombination. They allow the analysis of the outcome of indirect selection on forces that control genetic and phenotypic diversity. This approach is based on the assumption that genetic variation exists in these three rates of evolution; this has been demonstrated using heritability measurements and observed differences between sexes or closely related species (see, for example, 43-45 for the case of recombination modifiers). Recombination and mutation have been shown to be extremely variable across a species’ genome, with areas of low recombination or mutation, as well as hotspots of increased activity (46-47). The rate of migration and ability to disperse are typically heritable and vary between species and within environments for a single species (48-49).

We aim to better understand what are the similarities and discrepancies in the evolutionarily dynamics of these three evolutionarily important forces under similar patterns of environmental change. Elucidating the evolution of recombination, mutation and migration under fluctuating selection will lead to a more complete understanding of both fundamental evolutionary processes and the diverse pressures that have shaped the genome.

## Models of neutral modifiers in changing environments

Consider an infinite, randomly mating, haploid population. Individuals in this population are characterized by two types of biallelic loci: major loci, which control the phenotype and fitness of this individual, and modifier loci, which are selectively neutral. These neutral modifiers are assumed to control the force of interest, which can be the recombination rate between a pair of major loci, the mutation rate between alleles at the major loci, or the migration rate between two demes in a spatially subdivided population.

Intuitively, each of these phenomena, recombination, mutation and migration might be expected to increase genetic and phenotypic diversity and to enhance adaptation to environmental fluctuation that produces volatility in selection pressures. Under what conditions should evolution increase these rates and how do these conditions depend on temporal fluctuations in selection?

We address these questions by studying the evolution of the modifier loci and by determining the evolutionarily stable rates as functions of the pattern of fluctuation in selection experienced by the population. As in earlier analyses of neutral modifiers (50), we frame the problem in terms of the local stability of an equilibrium with one allele fixed at the modifier locus to invasion by another allele introduced at a low frequency near this equilibrium. The case where a stable multi-allele polymorphism exists at the modifier locus and a new modifier allele arises near this equilibrium (51), and the selection regime fluctuates, will be treated elsewhere.

### Evolution of recombination

Each individual is defined by three biallelic loci: two major loci *A/a* and *B/b* control the fitness of the individual while a third locus *M/m* is a modifier locus that controls the recombination rate between the two major loci, but is otherwise selectively neutral.

We study the evolution of the modifier locus *M/m* and determine the evolutionarily stable recombination rate as a function of the pattern of fluctuation in selection experienced by the population. This entails analysis of the stability of an equilibrium with only *M* present in the population to invasion by an allele *m*, introduced near this equilibrium.

To that end, we track the frequencies of the eight genotypes *MAB*, *MAb*, *MaB*, *Mab*, *mAB*, *mAb*, *maB*, *mab*. At each generation, the population experiences random mating, recombination, and selection, in that order.

There are three possible recombination rates depending on the mating type at the modifier locus: *MM*, *Mm*, and *mm* produce recombination rates *r*_1_, *r*_2_ and *r*_3_, respectively. With the three loci ordered as above and the modifier locus located on one side of the two major loci, let *R* be the recombination rate between the modifier locus and the two major loci. We assume no interference between recombination events occurring in the two intervals separating the two major loci and between the modifier locus and the nearest major locus.

Assume two possible types of selection regimes *T*_1_ and *T*_2_, such that the fitnesses, irrespective of the genotype at the *M/m* locus, can be represented as follows:

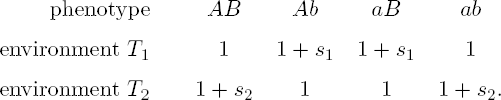

Thus if *s*_1_, *s*_2_ > 0, the genotypes that are better in one environment are less fit in the other. In this model, increased recombination should be favored since recombinant offspring are more fit than the non-recombinant ones (also see 14, 18, 22, 23).

### Evolution of mutation

The mutation model recapitulates that of Salathe et al. (29) and Liberman et al. (53), and we include it here for the purpose of comparison with recombination and migration. Each individual is defined by two biallelic loci: a major locus *A/a* controls the fitness of the individual while a second locus *M/m* is a modifier locus that controls the mutation rate between alleles at the major locus, but is otherwise selectively neutral.

We study the evolution of the modifier locus *M/m* and determine the evolutionarily stable mutation rate as a function of the pattern of fluctuation in selection. There are two possible mutation rates depending on the allele at the modifier locus: *M* and *m* produce mutation rates *μ*_1_ and *μ*_2_, respectively. Assume two possible types of selection regimes *T*_1_ and *T*_2_, such that the fitnesses are as follows:

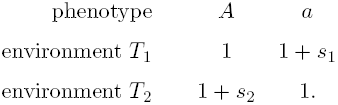

Assuming *s*_1_, *s*_2_ > 0, alleles that are better in one environment are less fit in the other. At each generation, the population experiences random mating, mutation, and selection, in that order.

### Evolution of migration

Here the population is divided into two demes *E*_*x*_ and *E*_*y*_, with different selection regimes. Individuals are characterized by two biallelic loci, a major locus *A/a* and a modifier locus *M/m*, where the major locus controls the fitness of an individual, while the modifier locus is assumed to be selectively neutral and controls the migration rate between the two demes. We investigate the evolutionary dynamics of the migration rate between *E*_*x*_ and *E*_*y*_ using an explicit population genetic model to track the frequencies of the four genotypes *AM*, *Am*, *aM* and *am*. At each generation, there is recombination and selection in each deme separately, after which individuals may migrate between the two demes. Again, we frame the question in terms of the local stability of the fixation equilibrium with only *M* present in the population, producing migration rate *ν*_*M*_, to invasion by allele *m*, which produces migration rate *ν*_*m*_. We assume *ν*_*m*_ and *ν*_*M*_ to be the same from *E*_*x*_ to *E*_*y*_ and from *E*_*y*_ to *E*_*x*_.

Within each deme, the selection regime varies temporally, with two possible environmental states, *T*_1_ and *T*_2_. In *T*_1_, the fitnesses are assumed to be

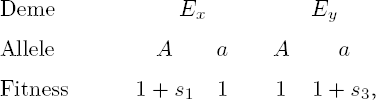

while in *T*_2_, they are

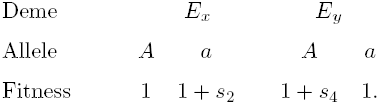

Thus, if *s*_1_, *s*_2_, *s*_3_, *s*_4_ > 0 within each deme, allele *A* is favored in one temporal state and allele *a* is favored in the other. This is an important assumption if the fitness regimes *T*_1_ and *T*_2_ covary between the two demes. Otherwise it is not important which environment is denoted as *T*_1_ or *T*_2_ and all that matters is that environmental change occurs independently within the two demes.

### Constant environment model as a reference

We first review models where selection is constant in time; this will serve as our reference model. Intuitively, in each of the three models, there exists one best-adapted genotype for a fixed environment, and the forces of recombination, mutation and migration act to reintroduce the less fit genotype that selection removes in each generation. Therefore, it is expected that the evolutionarily stable rates all evolve towards zero. This is indeed what we find in the mathematical analysis described in the **Supplementary Material**. This result is a haploid version of the reduction principle of Feldman and Liberman (51) (see also 30, 54).

### Changing environments: the general simulation model

Initially the population is fixed on allele *M* at the modifier locus, which determines a rate sampled randomly between 0 and 1 for modifiers of mutation and migration and between 0 and 0.5 for modifiers of recombination, and held constant thereafter. This population evolves for 1000 generations or until it reaches a stable polymorphic equilibrium at the phenotypic locus/loci. We then introduce allele *m* at the modifier locus at a small frequency (10^−4^) near this equilibrium. The new rate determined by *m* is chosen as the product of the resident rate of *M* and a number generated from an exponential distribution with mean 1. This allows us to test invading rates that are more often close to the resident rate, as well as those that are far from it. After 5000 generations, we determine whether the newly introduced allele was able to invade in the population; if the frequency of *m* is larger than its initial frequency of 10^−4^, we classify the outcome as an invasion. In this case, we expect the new allele to have had a selective advantage over the resident, and that it would reach fixation if the population continued to evolve. This is the initial invasion trial. If allele *m* did invade, the next invasion trial begins with this invading rate as the resident rate. If there was no invasion, the resident allele determines the same resident rate as in the previous trial. We then repeat the invasion steps described above. After at least 500 invasion trials and after the resident rate cannot be invaded in 50 consecutive trials, the final rate in the population is declared to be the evolutionarily stable rate.

To study the periodic case, the selection regime changes deterministically every *n* generations, making the environmental period 2*n*. To incorporate random temporal variation, we draw the waiting times between environmental changes from a gamma distribution. As a proxy for environmental variability we use the parameter *ψ*, which is the variance of the gamma distribution divided by the square of its mean. An exponential waiting time between environmental changes is a special case of the gamma distribution with variance equal to the mean squared, in which case *ψ* = 1. For simulations in which the variability parameter *ψ* is strictly larger than zero (non-periodic case), the final stable rate is computed as the average of the stable rates obtained in 10 different runs of the simulation.

## Analytic Results

Analytic, closed-form solutions for previous models of the evolution of modifier genes with temporal heterogeneity in selection have mostly been obtained when the environments change periodically, with short environmental runs. In the case of the evolution of mutation rates, Liberman et al. (53) showed that for period 2 and symmetric selection, increased mutation rates are always favored, and the mean fitness at equilibrium is an increasing function of the mutation rate. They also proved that with period 4, the stable mutation rate is 1/2. Moreover, the critical points of the mean fitness with respect to the mutation rate are the same mutation rates that cannot be invaded. Carja et al. (55) showed that these results hold in a model with spatial heterogeneity in selection pressure. Furthermore, for higher environmental periods, the parameter that controls the evolution is *m*_*b*_ = *m* + *μ* − 2*mμ*, where *m* is the migration rate and *μ* is the mutation rate, and the stable mutation rate is such that this parameter *m*_*b*_ is inversely proportional to the environmental period. Here, using the same mathematical tools, we derive analytical results for the evolutionary stable recombination and migration rates as functions of the environmental period, i.e. the length of the period of the fluctuating selection.

A general environmental cycle consists of *τ*_1_ + *τ*_2_ selection steps, the first *τ*_1_ of type *T*_1_ selection followed by *τ*_2_ of type *T*_2_ selection. We are able to derive exact analytical conditions for the local stability of fixation in *M* to invasion by *m* in the case *τ*_1_ = *τ*_2_ = 1, i.e. the environment changes every generation under the assumption that all selection coefficients in each of he three models are the same and equal to *s*. The mathematical analyses for modification of migration and recombination are presented in **Supplementary Material 1** and **2**. As noted above, in a temporally constant environment the reduction principle holds and smaller rates will always invade. On the other hand, with fluctuating selection of period 2, increased rates invade and the reverse of the reduction principle holds: a higher rate will invade a population fixed on a given resident rate. The evolutionarily stable rate is 1 for migration and mutation and 1/2 for recombination (the biologically feasible maximum). Note that our results in the haploid case are different from those for a modifier of recombination in a diploid model with period 2 described by Charlesworth (14).

We were able to obtain closed-form analytical solutions only under these very rigid symmetry assumptions and for environmental period of two. In order to understand the evolution of the three rates under more general models, we use simulations to determine the evolutionarily stable rate.

## Simulation Results

### Periodic environments

If the environment changes every generation, we confirm the result found in our mathematical analysis in the **Supplementary Material**. If the temporal environment changes deterministically every *n* generations (i.e. period of 2*n*) in both demes, the uninvadable rates are all decreasing functions of *n* (**Figure 1**). In all our results, the stable rates of mutation and migration are quantitively similar, suggesting that these two mechanisms lead to similar evolutionary dynamics. While the evolutionarily stable rates of mutation and migration are on the order of 1*/n*, in accord with previous analyses (see 26-28, 53, for modifiers of mutation), the evolutionarily stable recombination rate follows the same qualitative pattern, but does not appear to be related to 1*/n*, the rate of environmental change. **Figure 1** presents results for environmental rates *n* of *n* = 20 and above, while results for smaller environmental periods are shown in **Supplementary Figure S1**, where important differences in the three stable rates for small environmental periods can be seen. While the stable rate of recombination reaches the allowed maximum for all *n* < 6, mutation and migration exhibit different dynamics for small odd and even *n*. The maximal allowed rate is reached at *n* = 1 and *n* = 3 for migration, and *n* equal to 1,3,5, and 7 for mutation. This difference between odd and even *n* in the case of mutation modification was also observed by Liberman et al. (53). **Supplementary Figure S2** shows that these results are robust to different values of the selection coefficient *s*.

**Figure 1.**
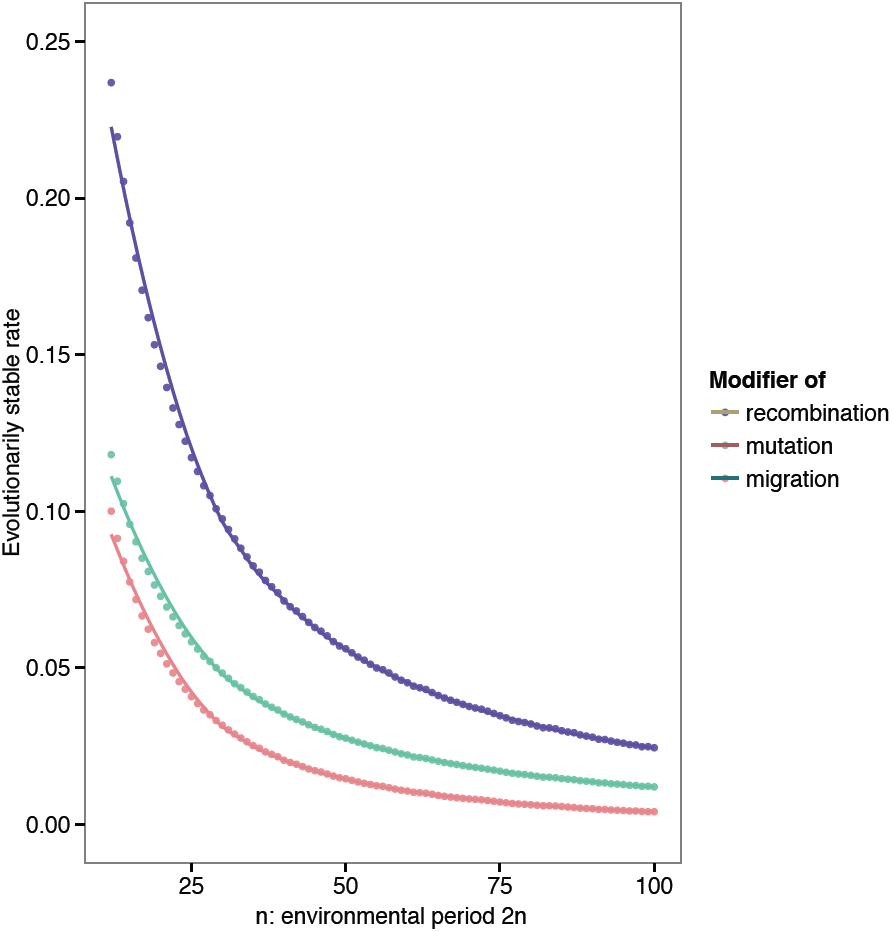
Evolutionarily stable rates as function of the number of generations before an environmental change *n*; symmetric selection and periodic environmental change. All selection coefficients are equal to 0.1. Recombination rates between modifier and major loci are 0 (*R* = 0 for recombination modification and *r* = 0 for mutation and migration modification). The rate of environmental change, *n*, equal to half of the environmental period is on the *x-axis*. The curves represent a fit to the data using a generalized additive model with penalized cubic regression splines.

This uninvadable rate also maximizes mean fitness at equilibrium. We show this in **Supplementary Figure S3** for recombination modifiers, and in **Supplementary Material 1** for migration modifiers; this was also found by Liberman et al. (53) for the evolution of mutation rates. Although this mean fitness principle holds for migration and mutation modifiers in diploid modifier models, it does not hold for the modification of recombination (56).

As recombination between the modifier locus and the gene(s) under selection increases, the strength of secondary selection on the modifier decreases because the linkage disequilibrium between these genes decreases. **Supplementary Figure S4** shows that the stable evolutionary rates decline with increasing recombination rate between the modifier and the major locus/loci. In earlier studies of neutral modifiers of mutation and recombination, the induced selection on the modifier locus was of the order of the square of the disequilibrium between the major and the modifier loci (54). Even if this effect is exaggerated somewhat by the environmental fluctuations, it remains weaker than the effect of environmental volatility.

### Environmental variability

When the waiting times between temporal changes are random, the 1*/n* rule no longer holds. **Figure 2** shows the stable rates of recombination, mutation and migration when the expected time before an environmental change is 10 generations and this temporal change is sampled from a gamma distribution with variability parameter *ψ* represented on the *x-axis*. Environmental variability *ψ* = 0 is the case of two periodically changing environments with period 10 and recaptures the behavior seen in **Supplementary Figure S1**. Non-zero variability *ψ* drastically changes the behavior of the system. We find that the evolutionarily stable rates can be up to two orders of magnitude lower than in the periodic regimes and depend strongly on the selection coefficients; in fact, the decrease in the stable rate is steeper as selection becomes weaker.

**Figure 2.**
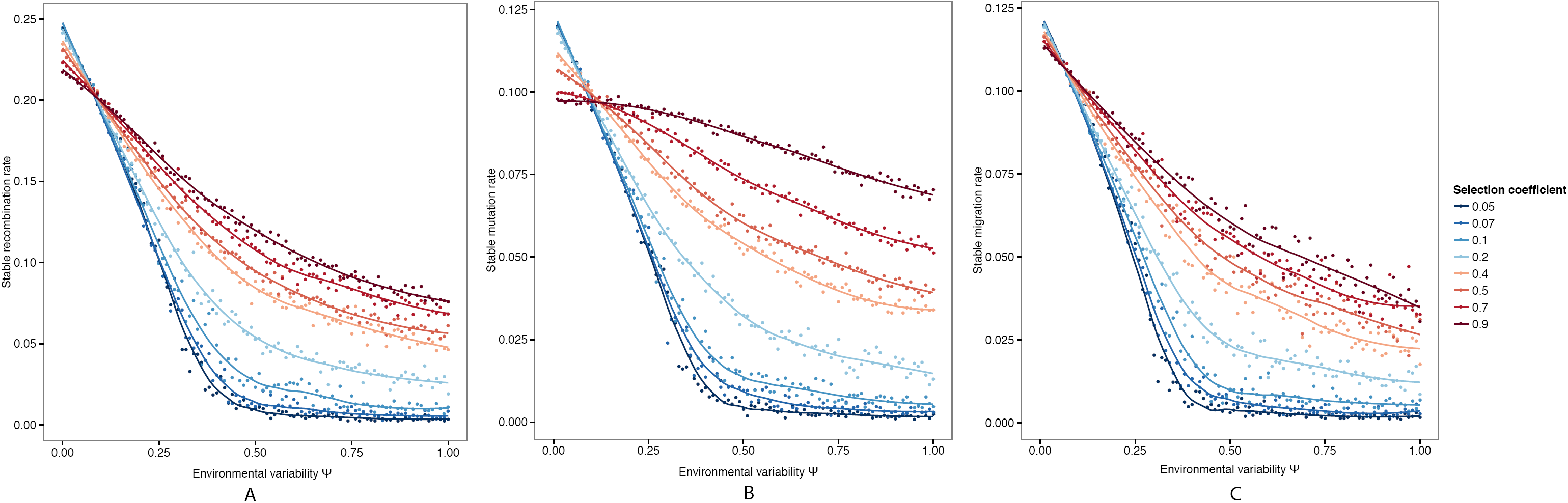
Evolutionarily stable rates as function of the environmental variability *ψ*; symmetric selection and random environmental change. All selection coefficients are equal and as presented in the legend. Recombination rate between modifier and major loci is 0.1. The variability parameter *ψ* is presented on the *x-axis*. The curves represent a fit to the data using a generalized additive model with penalized cubic regression splines. The three panels present the results for evolutionarily stable rates of recombination in **Panel A**, mutation in **Panel B** and migration in **Panel C**.

### Asymmetric selection

When the fitnesses of the genotypes depend on the environment, we observe a threshold phenomenon where, if the selection coefficients are similar enough, the evolved rates are the same as in symmetric landscapes (i.e. with a single selection coefficient *s*); however, past a certain threshold difference in selection coefficients, there is a sudden drop in the evolutionarily stable rate and non-zero rates can no longer be stable. The threshold substantially limits the circumstances under which non-zero recombination, mutation and migration rates can be maintained as shown in **Figure 3, Panels A and B** for both periodic and random environmental changes. This was observed for mutation modifiers (see 29, 53) and is robust to differences in environmental variability ψ and environmental mean waiting times (see **Supplementary Figures S6 and S7**).

**Figure 3.**
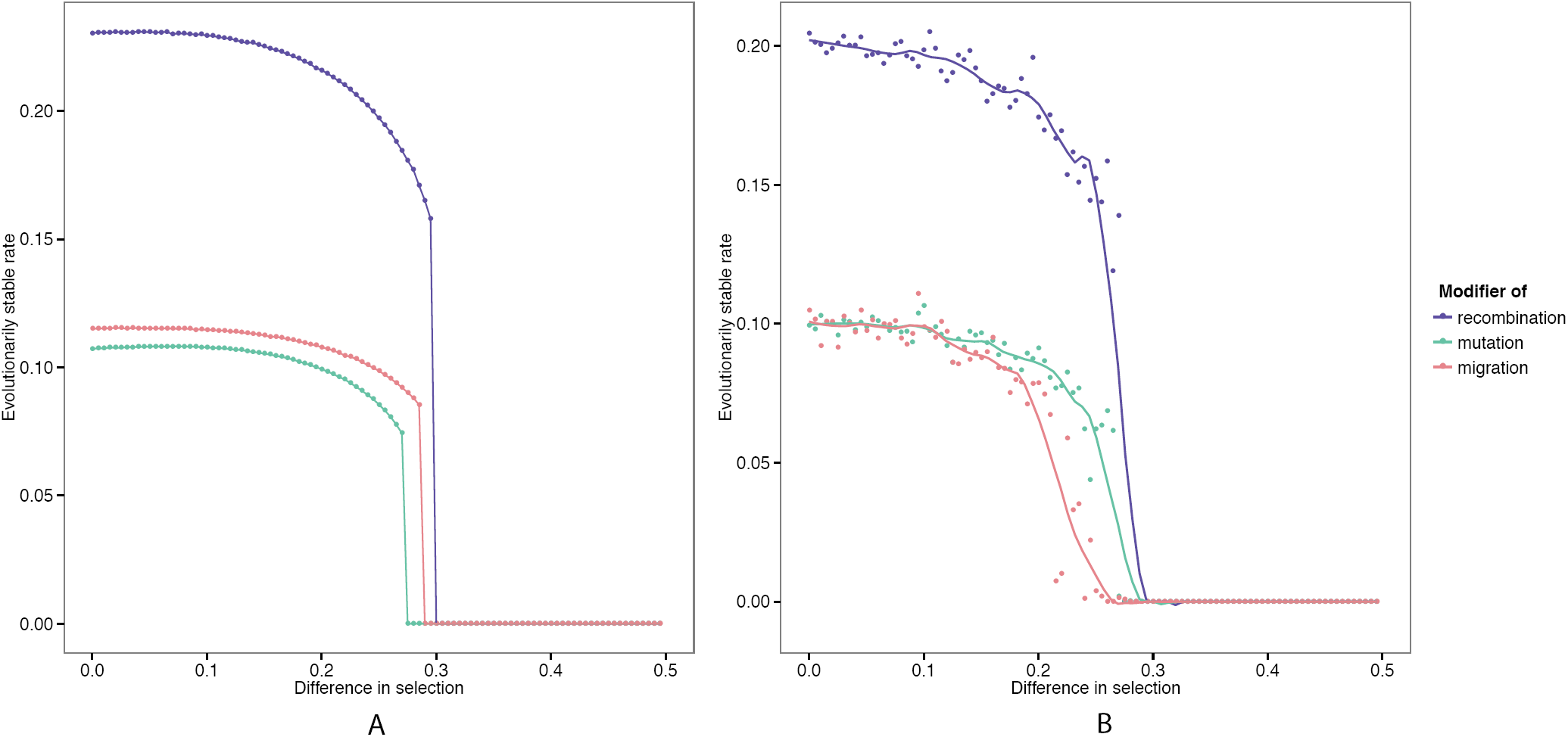
Evolutionarily stable rates as function of the number of generations before an environmental change *n*; asymmetric selection. For recombination and mutation: *s*_2_ = 0.5 and the difference in selection *s*_2_ – *s*_1_ is on the *x-axis*. For migration: *s*_2_ = *s*_4_ = 0.5 and *s*_1_ = *s*_3_ with the difference in selection *s*_2_ – *s*_1_ on the *x-axis*. Recombination rates are *R* = 0.1 for recombination modification and *r* = 0.1 for mutation and migration modification. **Panel A**: Periodic environment, rate 10 (period 20). **Panel B**: Mean waiting time between environmental changes is equal to 10 and variability parameter *ψ* = 0.1. The dots represent averages over 10 runs of the simulation. The curves represent a fit to the data using a generalized additive model with penalized cubic regression splines.

## Discussion

A wide array of mechanisms may increase genetic diversity in response to fluctuation in selection pressures. Here we explore the dynamics of three important evolutionary forces in temporally fluctuating environments: recombination, mutation and migration. To this end, we use deterministic, haploid population genetic, modifier models to determine the uninvadable evolutionary rates for these three different mechanisms. An important parameter in our analysis is the rate of environmental change, and we explore both periodic and random waiting times before a change in the selection regime.

These models track the evolution of a modifier linked to a locus under fluctuating selection and display interesting similarities in the evolutionary dynamics of mutation, recombination and migration. These similarities may stem from the fact that all three forces can enhance a population’s adaptation to a changing environment by increasing the genetic diversity on which selection can act. In a sense, they all allow for anticipation of the pattern of selection change but they all also create a load. In that context, mutation, recombination and migration can be considered bet-hedging strategies. However, the type of selection acting in the these models is different from what is classically called bet-hedging because we consider an infinite population size model with no demography (6).

For each of these forces, our models reproduce and extend some of the qualitative messages of previous theoretical work. In constant environments, the reduction principle holds and the rates of mutation, migration and recombination all evolve towards zero. When environments change periodically through time, these rates can evolve to non-zero values, each decreasing with increasing environmental period. Previous work in the evolution of mutation rates (27-28) found that mutation evolves to be in synchrony with the rate of environmental change: order of 1/*n*, where 2*n* is the environmental period. Here we use a genetic model to show that the rate of migration between two demes with distinct selection pressures also follows this pattern and the evolutionarily stable rates of migration are quantitively similar to those of mutation. This is because mutation and migration both act to introduce the fitter alternative alleles into the changed environment. We also show that the rate of recombination follows the same qualitative pattern as the rates of mutation and migration and find the stable recombination rate as a function of the environmental period.

Previous work has mostly focused on periodic, symmetric environmental changes. The total genetic load differs between randomly changing environments and periodically fluctuating environments and therefore, the evolutionary outcome between random and periodically fluctuating selection regimes is expected to be different (57). These essentially different forces (recombination, mutation and migration) are surprisingly similar in their evolutionary response to variability in environmental changes; increase in the variability parameter *ψ* leads to a decrease in all three evolutionary rates. We show that an increase in the variability of the environmental conditions leads to a decrease in the forces of recombination, mutation and migration. We hypothesize that, as this variability increases, the longer stretches of constant selection that favor reduction shape the dynamics of the system. This decreased role of short environmental durations would explain why the optimal rates of these three forces are small when the environmental fluctuation is random. Moreover, the uninvadable rates increase with the selection intensity; when selection is weak, the evolutionary dynamics track environmental shifts slowly and the occasional very long runs of the same environment pose a disadvantage to a high recombination, mutation or migration rate. As the selection pressure increases though, the system is better able to respond to shorter runs and shifts in the environment. As a consequence, the stable rates increase so that the average environmental duration decides the evolutionary fate of these three forces.

The question of optimal rates of genomic evolution needs to be further explored under more general selection scenarios driven by both temporal and spatial change. An important limitation of our models is the assumption of infinite population size: in finite populations under a constant environment, drift may be important in shaping the stable rates of these three forces we have studied here (58-60). A better understanding of the interplay of drift and environmental change in the evolution of recombination, mutation and migration is needed in both experimental and theoretical settings, since we expect that effective population size and genetic drift could play important roles in these systems. Previous work on evolution of migration rates / dispersal showed that, with drift, migration will be selected as a way to avoid kin competition (61). Similarly, random genetic drift caused by sampling in finite populations can affect the evolutionarily stable recombination rate: drift can generate negative disequilibria among selected loci (63) such that beneficial alleles at one locus become associated with deleterious alleles at other loci. Modifiers that increase recombination could then spread in the population because they help regenerate good combinations of alleles and will rise in frequency along with these alleles (59-60, 62). In large populations, it has been proposed that selection on recombination should be inversely proportional to population size (64). However, even with large numbers of individuals, a drift based-advantage to recombination can occur as long as there is spatial structure (63) or selection acts on a large number of loci (64).

Another limitation of our model is the assumption of a small number of loci on which selection acts rather than many loci under selection, controlled by many modifiers; we expect that the qualitative patterns found here will hold in systems with more than two main loci, though the evolutionarily stable values of the three forces may change (20, 60).

A major theoretical challenge is to understand how the different forces discussed here combine to influence the evolution of genetic and phenotypic diversity. Here, we have analyzed each of these mechanisms separately; natural populations generally use a combination of the three forces to counteract the stresses induced by a change in selection. An understanding of the interplay between the different types of diversity introduced by these three different processes is needed in theoretical population genetics. For example, it has been shown that, with migration among populations, spatial heterogeneity in selection can generate positive or negative linkage disequilibrium and select for recombination even in the absence of genetic drift or temporal heterogeneity (17). This is because natural selection varying across space maintains local differences in gene frequencies, and with migration these differences in frequency can generate linkage disequilibria within a deme and create a selective pressure for increased recombination. In fact, including migration extends the range of epistasis over which recombination can be favored (17). In our recent study of mutation modification in two populations connected by migration we observed that the parameter of interest that controls the evolution of the system is a non-linear function of the mutation and migration rates and its evolutionarily stable rate is inversely proportional to the environmental period (55). Mutation and recombination have also been studied together in models for evolution of recombination in constant environments; if the major loci are at an equilibrium between selection against deleterious alleles and mutation towards them, recombination can increase in the population if the linkage disequilibrium is negative (15, 21).

Overall, we find that these three, essentially different, processes and mechanisms are surprisingly similar in how they respond to changes in selection pressure, as each of them constitutes a form of genetic bet-hedging and endows a population with the genetic diversity that can accommodate the changes in their selective environments. Our models show that knowing the difference in selection pressure between environmental regimes is not sufficient for predicting the long-term advantage of these three forces of evolutionary change. Knowledge of the duration, shape and randomness of the environmental regimes is also essential. Similarly, knowing how the environment fluctuates is of little help without knowing the extent of asymmetry in environmental pressure. In a broader perspective, we believe that future work needs to consider the interactions across multiple dimensions in the parameter space in order to increase our understanding of the evolution of recombination, migration and mutation, as well as other traits that contribute to adaptation in a changing world.

## List Of Supplementary Figures

**Figure S2.**
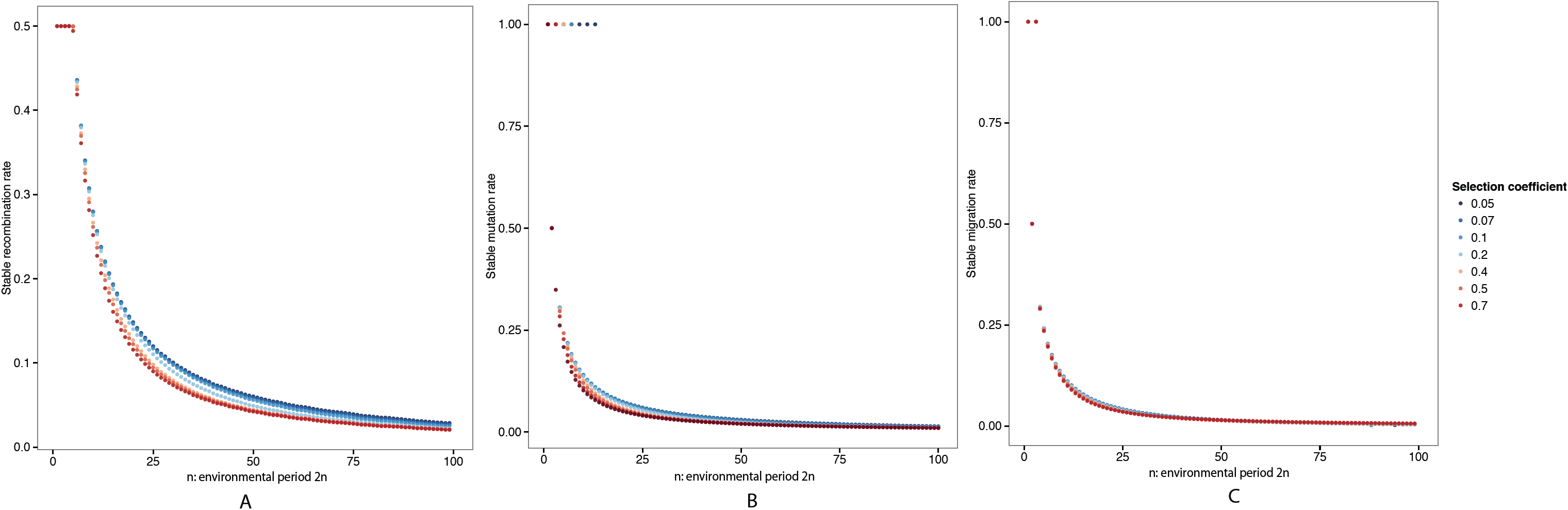
Supplementary Figure S2. Robustness of Figure 1 to different symmetric selection pressures. All selection coefficients are equal and in the legend. Recombination rates between modifier and major loci are *R* = *r* = 0. The rate of environmental change *n*, equal to half of the environmental period, is on the *x-axis*. The curves represent a fit to the data using a generalized additive model with penalized cubic regression splines.

**Figure S3.**
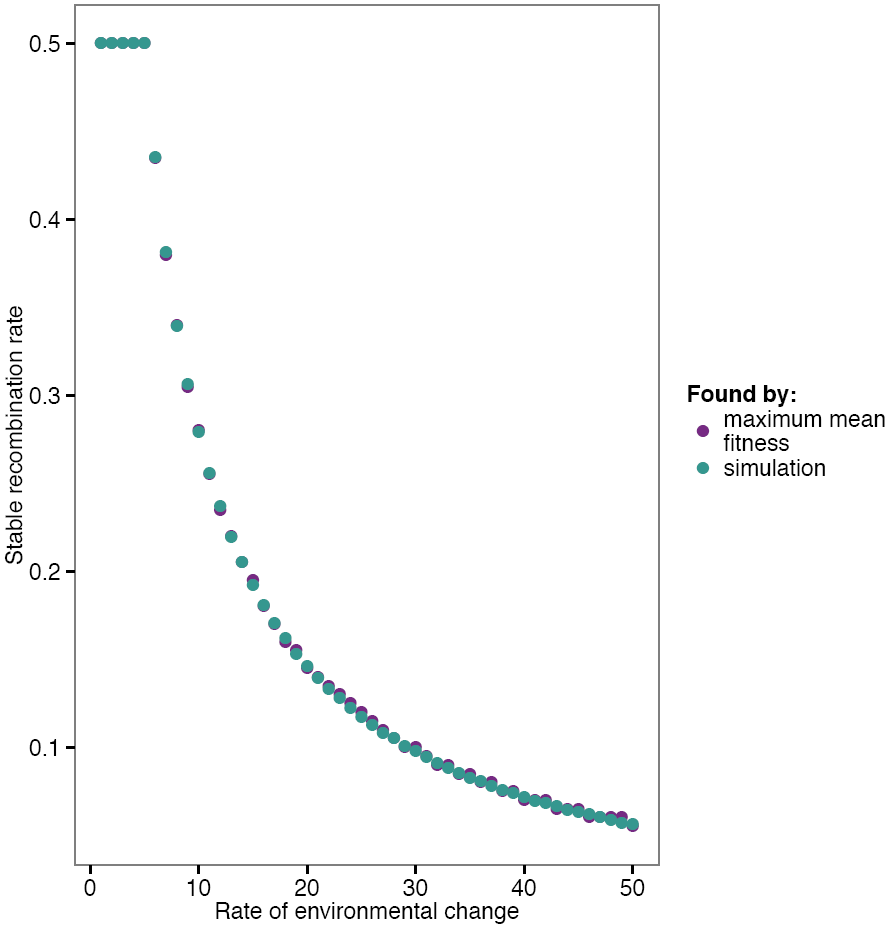
Supplementary Figure S3. The stable recombination rate and the maximum mean fitness at equilibrium. All selection coefficients are equal to 0.1. Recombination rates between modifier and major loci are *R* = *r* = 0. The rate of environmental change *n*, equal to half of the environmental period, is on the *x-axis*. The rates obtained by maximizing mean fitness and the simulation results are equal to 2 decimal points.

**Figure S4.**
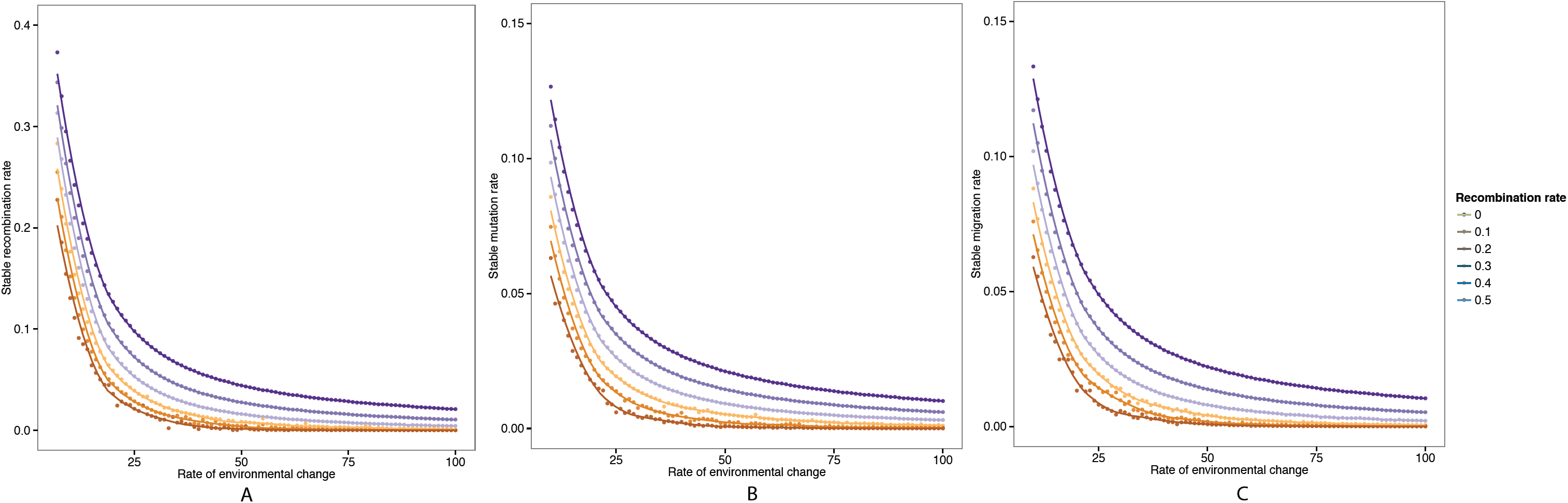
Supplementary Figure S4. The role of recombination between major loci and modifier (R for recombination modification and r for mutation and migration modification). Symmetric selection coefficient is *s* = 0.4. Recombination rate as in the legend. The environment changes periodically, with rate *n* on the *x-axis*. The curves represent a fit to the data using a generalized additive model with penalized cubic regression splines.

**Figure S1.**
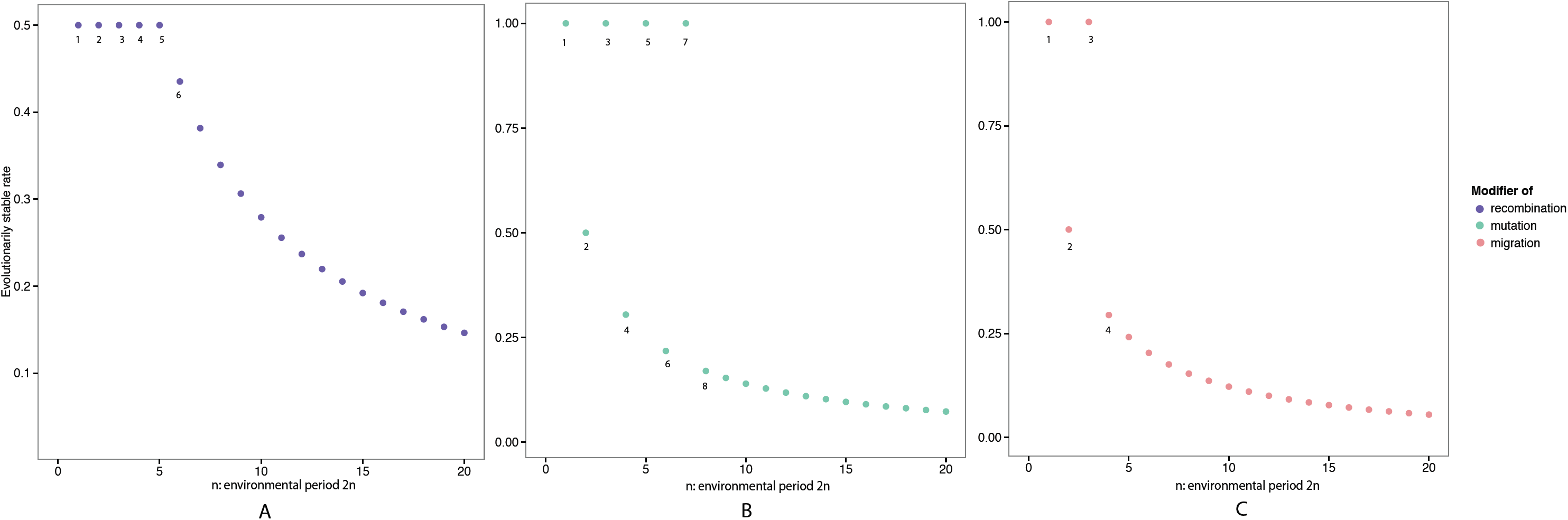
Supplementary Figure S1. The same as Figure 1, smaller periods. All selection coefficients are equal to 0.1. Recombination rates between modifier and major loci are *R* = *r* = 0. The rate of environmental change *n*, equal to half of the environmental period, is presented on the *x-axis*. The curves represent a fit to the data using a generalized additive model with penalized cubic regression splines.

**Figure S5.**
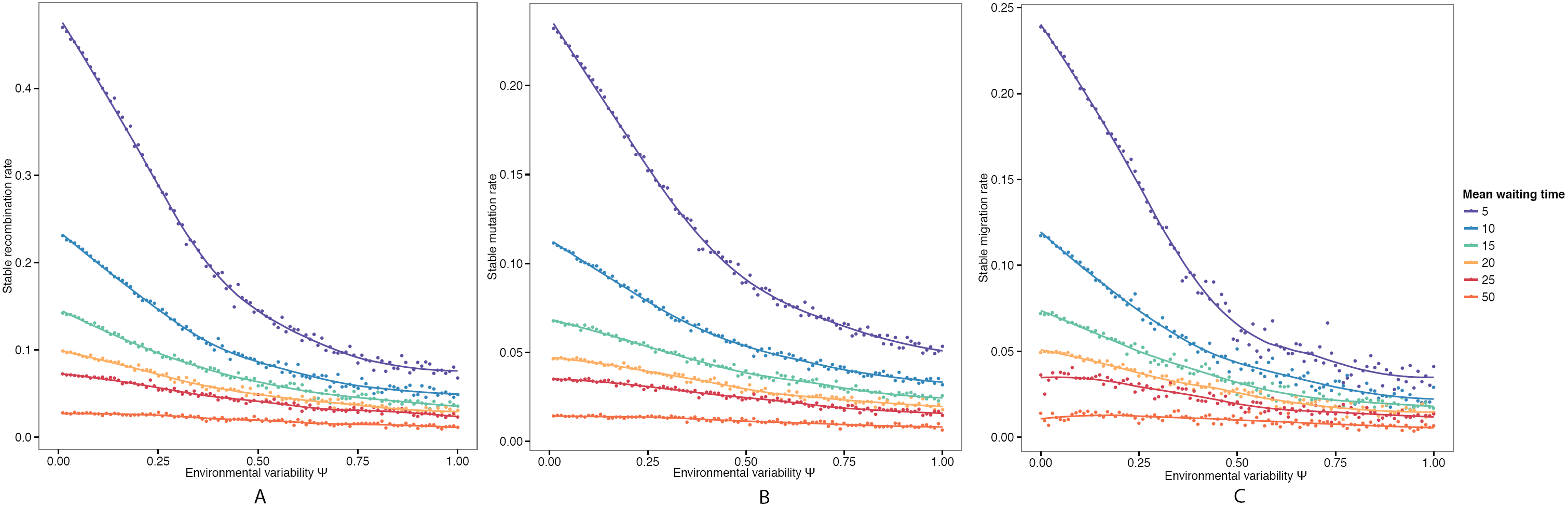
Supplementary Figure S5. Robustness of Figure 2 to different environmental mean waiting times. Symmetric selection coefficient is *s* = 0.4. Recombination rates are *R* = 0.1 and *r* = 0.1. Mean waiting time between environmental changes is in the legend. The variability parameter *ψ* is presented on the *x-axis*. The dots represent the average over 10 runs of the simulation. The curves represent a fit to the data using a generalized additive model with penalized cubic regression splines.

**Figure S6.**
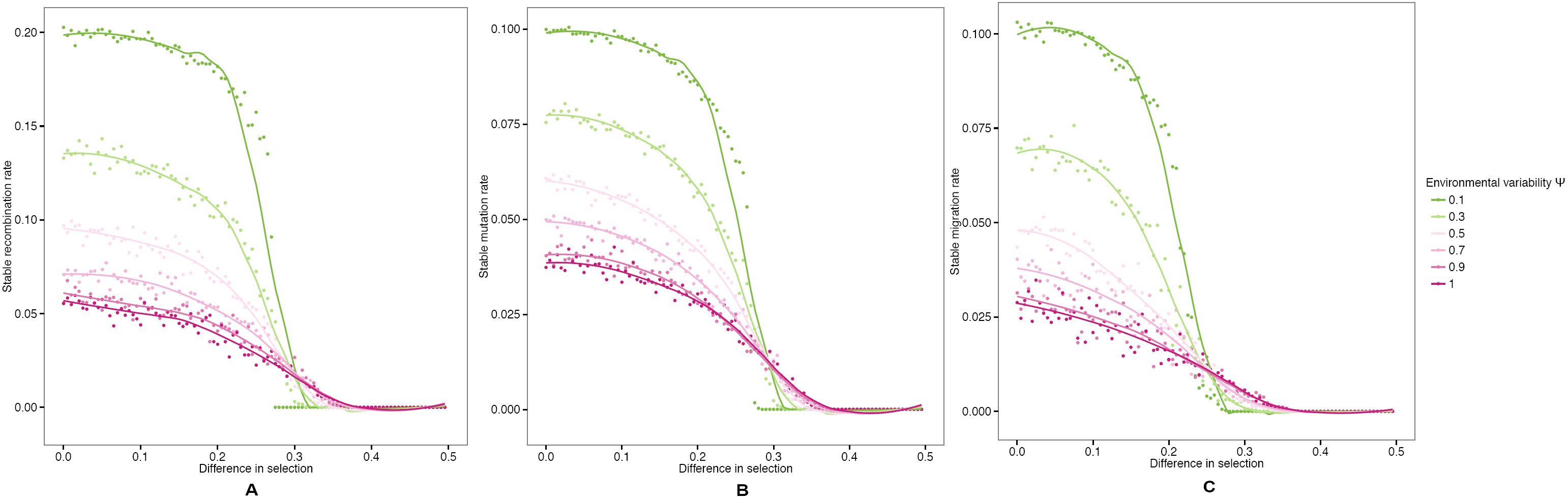
Supplementary Figure S6. Robustness of Figure 3 to different environmental variabilities *ψ*. For recombination and mutation: *s*_2_ = 0.5 and the difference in selection *s*_2_ — *s*_1_ is on the *x-axis*. For migration: *s*_2_ = *s*_4_ = 0.5 and *s*_1_ = *s*_3_ with the difference in selection *s*_2_ — *s*_1_ on the *x-axis*. Recombination rates are *R* = 0.1 and *r* = 0.1. Mean waiting time between environmental changes is 10. The dots represent the average over 10 runs of the simulation. The curves represent a fit to the data using a generalized additive model with penalized cubic regression splines.

**Figure S7.**
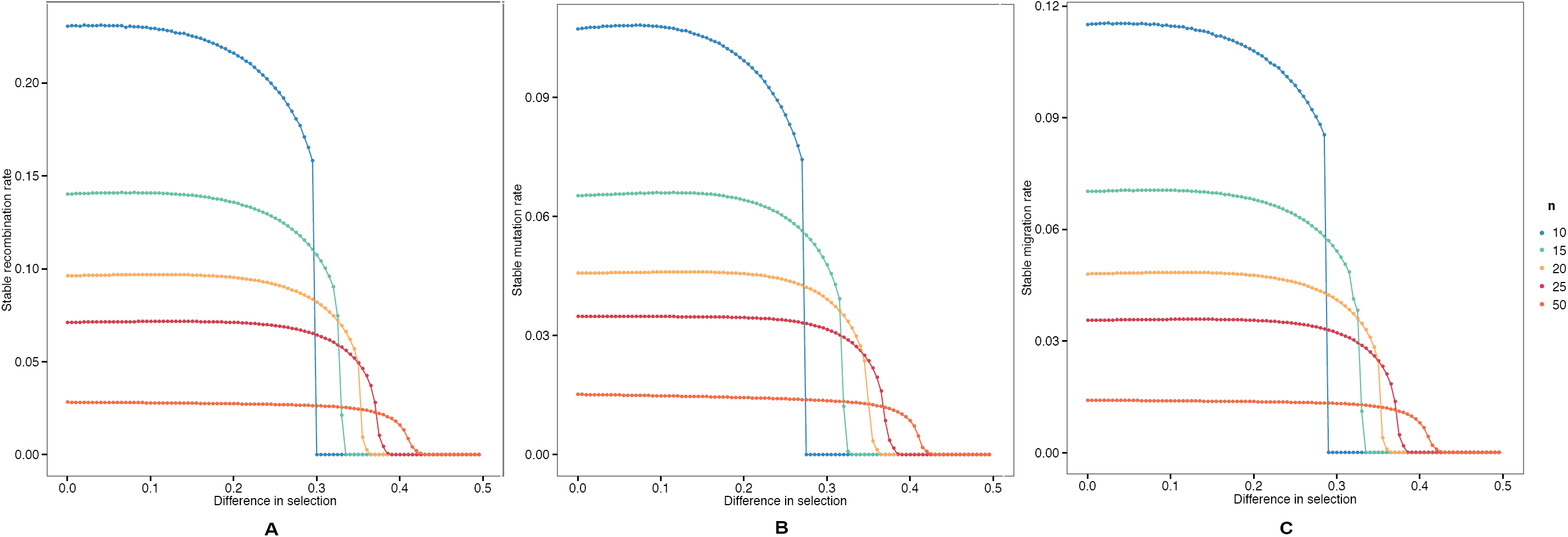
Supplementary Figure S7. Robustness of Figure 3 to different number of generations before an environmental change *n*. For recombination and mutation: *s*_2_ = 0.5 and the difference in selection *s*_2_ — *s*_1_ is on the *x-axis*. For migration: *s*_2_ = *s*_4_ = 0.5 and *s*_1_ = *s*_3_ with the difference in selection *s*_2_ — *s*_1_ on the *x-axis*. Recombination rates are *R* = 0.1 and *r* = 0.1. Environmental variability *ψ* = 0. The dots represent averages over 10 runs of the simulation. The curves represent a fit to the data using a generalized additive model with penalized cubic regression splines.

